# Notch signaling determines cell-fate specification of the two main types of vomeronasal neurons of rodents

**DOI:** 10.1101/2021.10.26.466003

**Authors:** Raghu Ram Katreddi, Ed Zandro M. Taroc, Sawyer M Hicks, Jennifer M Lin, Shuting Liu, Mengqing Xiang, Paolo E. Forni

**Affiliations:** Department of Biological Sciences, University at Albany, State University of New York, Albany, NY, United States; The RNA Institute, University at Albany, State University of New York, Albany, NY, United States; The Center for Neuroscience Research, University at Albany, State University of New York, Albany, NY, United States; State Key Laboratory of Ophthalmology, Zhongshan Ophthalmic Center, Sun Yat-sen University, Guangzhou 510060, China

**Keywords:** Single cell sequencing, Vomeronasal Organ, Notch signaling, Dll4, neuronal dichotomy, mouse, neuronal differentiation

## Abstract

The ability of terrestrial vertebrates to find food, mating partners and to avoid predators heavily relies on the detection of chemosensory information from the environment. The olfactory system of most vertebrate species comprises two distinct chemosensory systems usually referred to as the main and the accessory olfactory system. Olfactory sensory neurons of the main olfactory epithelium detect and transmit odor information to main olfactory bulb (MOB), while the chemosensory neurons of the vomeronasal organ detect semiochemicals responsible for social and sexual behaviors and transmit information to the accessory olfactory bulb (AOB). The vomeronasal sensory epithelium (VNE) of most mammalian species contains uniform vomeronasal (VN) system with vomeronasal sensory neurons (VSNs) expressing vomeronasal receptors of the V1R family. However, rodents and some marsupials have developed a more complex binary VN system, where VNO containing a second main type of VSNs expressing vomeronasal receptors of the V2R family is identified. In mice, V1R and V2R VSNs form from a common pool of progenitors but have distinct differentiation programs. As they mature, they segregate in different regions of the VNE and connect with different parts of the AOB. How these two main types of VSNs are formed has never been addressed. In this study, using single cell RNA sequencing data, we identified differential expression of Notch1 receptor and Dll4 ligand among the neuronal precursors at the VSN dichotomy. We further demonstrated with loss of function (LOF) and gain of function (GOF) studies that Dll4-Notch1 signaling plays a crucial role in triggering the binary dichotomy between the two main types of VSNs in mice.

**Graphical Abstract:** 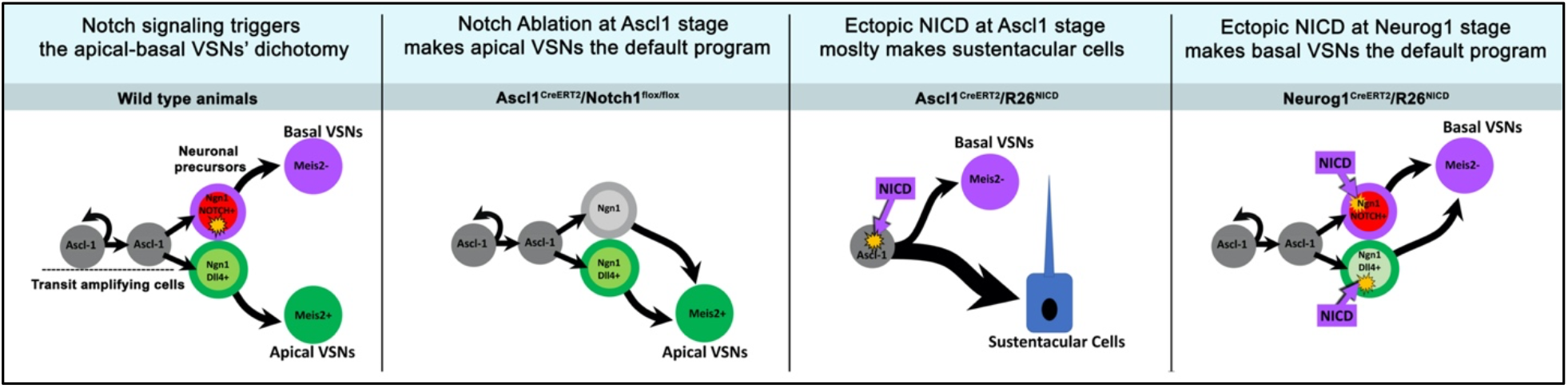

## Introduction

Neural stem/progenitor cells can give rise to multiple neuronal cell types that differ in gene expression, functions, and neuronal connectivity. Investigating the molecular mechanisms that establish different neuronal cell-fates is crucial to understand how neuronal systems evolve, form and, to identify molecular mechanisms underlying neurodevelopmental disorders [1-3].

The vomeronasal organ (VNO) is a specialized chemosensory organ that, in many vertebrate species, is located at the base of the nasal cavity [4]. The VNO is responsible for the detection of semiochemicals, molecules that can trigger stereotypical mating/sex behaviors, parental behaviors, and predator avoidance [5-8]. Most of the vertebrates having a functional VNO, have neurons expressing vomeronasal receptor genes of the V1R family [9]. However, the vomeronasal sensory epithelium (VNE) of mice, rats and opossum consists of two main types of vomeronasal neurons [10]. VSNs that express Gαi2 G protein subunit, receptors of the V1R family and the transcription factor Meis2 are fairly conserved population across many vertebrate species [11]. The second population that has been reported in rodents and few other animal species is formed by the VSNs that express the Gαo G protein subunit, receptors of the V2R family and the transcription factor tfap2e (AP-2ε) [11-14]. In mice, the V1R expressing neurons are mostly distributed in the apical territories of VNE and for this reason they are often referred to as apical VSNs. Conversely, the V2R expressing neurons that, for the most part, are located in basal regions of the VNE and around the vasculature, are called basal VSNs [15]. Both apical and basal VSNs detect distinct types of ligands, connect to different areas of the accessory olfactory bulb and control distinct behaviors [16-19].

In mice, the VSN neurogenesis starts during embryonic development around E11.5 and continues, throughout the life, starting from a limited number of progenitors in the marginal zones of the VNO [15, 20-23]. How the cell fate of apical and basal VSN types is established has not been fully understood. A pivotal study by Enomoto and coworkers previously identified that the transcription factor Bcl11b plays a key role in controlling the correct establishment of apical and basal VSNs in the developing VNO [11]. However, what extrinsic signaling pathways control the expression of Bcl11b in the VNO has not been investigated until now. Moreover, the same study identified, the transcription factor AP-2ε as a potential target of Bcl11b and suggested a role for AP-2ε in controlling basal neuron differentiation. In a follow-up study, we proposed that the AP-2ε is not responsible for initiating the apical vs basal identity bifurcation but is crucial for the expression of several genes defining, and maintaining, the basal VSN’s molecular identity [14]. In fact, we observed that after AP-2ε loss of function, VSNs with basal identity lose the expression of basal genes (Gao and V2Rs) while they switch to expressing apical genes such as Gai2 and V1Rs [14].

In this study we adopted a single cell sequencing strategy to investigate the mechanisms involved in cell fate specification and differentiation of apical and basal VSNs. By following the transcriptomic profile of postnatal stem cells/progenitors and immature VSNs, we identified differential expression of Notch-1 receptor and Dll4 ligand across Neurog1/Neurod1 positive VSNs’ precursors. Following this hint, we adopted both loss of function (LOF) and gain of function (GOF) *in-vivo* strategies to test the role of this signaling pathway in establishing the VSNs’ binary differentiation. LOF and GOF strategies differentially affected the switch of apical vs basal VSN cell fate with Notch signaling LOF directing cells towards apical and GOF towards basal VSN fate. Our data on the VNO offer a new example of how Notch-mediated cell fate choice is a conserved strategy for establishing binary cellular diversification and increasing the neuronal repertoire in developing neuroepithelia.

## RESULTS

### Single cell profiling of the whole adult VNO identifies VSN dichotomy

In mice, adult neurogenesis is seen throughout the life at marginal zones of the vomeronasal epithelium [23, 24]. To identify multiple cell types like stem cells, VSNs and non-neuronal cell types, we performed single cell RNA sequencing (sc-RNA seq) from dissociated whole VNO. We used Seurat, R package, to filter out low quality cells and perform clustering and analyze sc-RNA-seq data [25]. A total of 10,582 single-cell transcriptomes passed the quality control measures. Based on the expression of top 2000 highly variable features across the population, cells were clustered in Seurat object1 and visualized using uniform manifold approximation projection (UMAP) [26] (Fig. 1A). We identified the neuronal dichotomy (circled region in Fig. 1A) and annotated the cell clusters that belong to stem cells, neuronal progenitors, immediate neuronal precursors and immature VSNs based on the following gene expression pattern: Sox2/Ascl1 – Neuronal progenitors; Neurog1/Neurod1 – Immediate neuronal precursors; Gap43-immature VSNs; Meis2-apical VSNs; Tfap2e-basal VSNs (Supplementary Fig. 1). We focused our further clustering and analysis only on Ascl1+, Neurod1+ and Gap43+ cells (circled in Fig. 1A) as our main interest is to characterize the mechanisms underlying the VSN cell fate specification. We subset these clusters into Seurat object2 (Fig. 1B) and re-clustered based on highly variable features among the selected cells. The expression pattern of known apical and basal specific genes as shown in the heatmap (Fig. 1B’’), further corroborated the sc-RNA seq analyzed data with the literature [11, 14, 20]. Interestingly, Sox2 expression was found to be extended from the Ascl1+ progenitors’ stage to Neurog1+ and to a certain extent of Neurod1+ neuronal precursors’ stage (Fig. 1B’, Fig. 2B), similar to what previously shown in the main olfactory epithelial studies [27, 28]. In addition, we found that Neurod1 expression, that starts along with Neurog1 after Ascl1, is retained beyond the dichotomy in both apical and basal VSN lineages (Fig. 1B’, Fig. 2B). We labelled Neurog1+/Neurod1+ cells before the dichotomy as immediate neuronal precursors, whereas Neurod1+ cells in apical and basal lineages, after the dichotomy, as putative apical and basal neuronal precursors respectively.

**Figure1:**
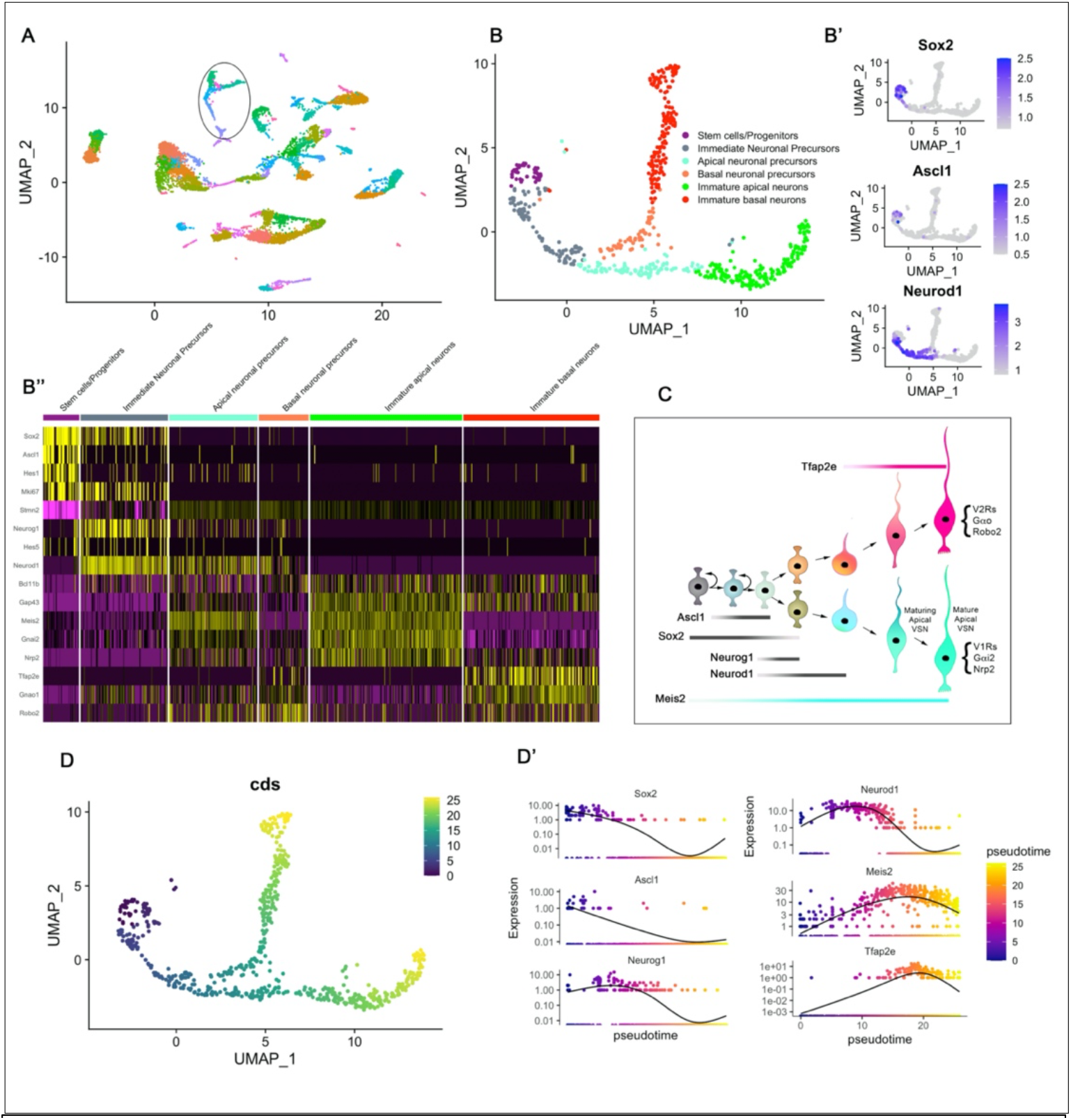
sc-RNA-seq identified adult neurogenesis and VSN apical-basal dichotomy in the mouse VNO. **A)** UMAP dimensional reduction plot of Seurat object1 shows neuronal and non-neuronal cell clusters of VNO. Each color corresponds to a cluster of cells that have similar transcriptomic profile. Clusters that are circled consists of neuronal progenitors, precursors and immature VSN apical and basal populations. **B)** Seurat object2 generated from highlighted clusters of Seurat objet1 (encircled) distinguishes VSN neuronal dichotomy and identifies cell clusters specific to neuronal progenitors, immediate neuronal precursors, apical and, basal neuronal precursors and apical and, basal immature VSNs. **B’)** Feature plots of Sox2, Ascl1 and Neurod1 in Seurat object2. **B’’)** Heatmap of the known genes that are specific to each cluster. **C)** Summary cartoon depicting VSN dichotomy from stem cell/ neuronal progenitor stage and dynamic transcriptomic expression of stage specific transcription factors. D) The single cell pseudotime trajectory of Seurat object2 predicted by Monocle Seurat wrapper and visualized by UMAP. Cells are ordered in pseudotime by choosing Ascl1+ cells as root node and colored in a gradient from purple to yellow. D’) Dynamic expression of small set of genes as a function of pseudotime.

**Figure 2:**
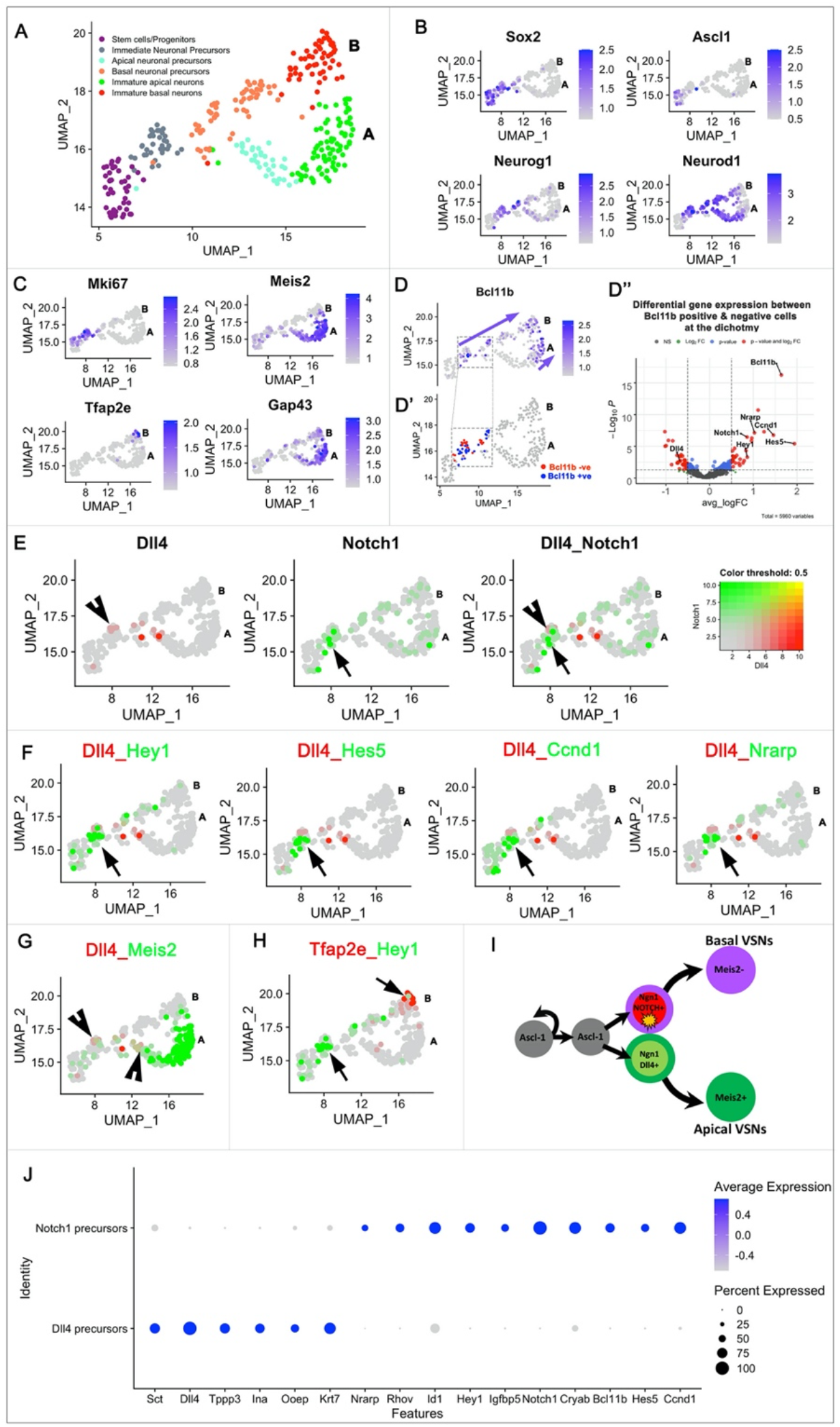
sc-RNA seq analysis identifies Notch1-Dll4 signaling in establishing VSN dichotomy. For all the plots in this figure panel, alphabets A and B denotes apical and basal VSN branches. **A)** UMAP dimension plot of Seurat object3 that is specifically focusing on the VSN dichotomy. **B, C)** Feature plots of known marker genes that are specific to neuronal progenitors, precursors and immature apical and basal VSNs. **D)** Feature plot of Bcl11b expression at the VSN dichotomy. Arrow mark highlights continuous gene expression in the basal VSN branch, whereas it appears at a later stage in the apical VSN branch. **D’)** Feature plot highlighting Bcl11b positive and negative cells at the dichotomy considered for differential gene expression analysis. **D’’)** Volcano plot highlighting Notch related genes differentially expressed between Bcl11b positive and negative clusters. **E)** Feature plots of Dll4 in red (arrowhead) and Notch1 in green (arrow) show their complimentary expression at the VSN dichotomy. **F)** Co-expression feature plots of Dll4 vs downstream Notch signaling targets like Hes5, Hey1, Nrarp and Ccnd1 (follow arrow marks) show that active Notch signaling happens transiently at the dichotomy and early stages of basal VSN trajectory. **G)** Co-expression feature plot of Dll4 vs Meis2 (arrowhead) shows Dll4 expression towards apical VSN branch **H)** Co-expression feature plot of Hey1 vs Tfap2e (arrows) shows active Notch signaling hapenning towards basal VSN branch. **I)** Summary schematic showing the summary of Notch1-Dll4 signaling required for establishing VSN dichotomy. **J)** Dot plot of a few selected genes that are differentially expressed between Dll4 and Notch1 positive cluster at the VSN dichotomy.

We further implemented Monocle single cell trajectory analysis on Seurat object2 [29, 30]. In this analysis, we chose Ascl1 positive neuronal progenitors as a root node and rest of the single cells in Seurat object2 were ordered based on pseudotime, which is a unit of progress. This revealed a branched trajectory confirming the VSN dichotomy obtained from Seurat analysis (Fig. 1D, D’).

### Differential gene expression identifies the expression of Notch signaling related genes at the VSN dichotomy

To further focus on the VSN dichotomy, we re-clustered Ascl1+ progenitors and VSNs’ neuronal precursors to form Seurat object 3 (Fig. 2A). Feature plots of Ascl1, Sox2, Neurog1 and Neurod1 (Fig. 2B) identified the temporal transcriptional cascade as neuronal progenitors proliferate and differentiate into immediate neuronal precursors. Further analysis showed, as expected [11, 14], a distinct expression of the transcription factor Meis2 in post-mitotic (Ki67 negative) cells acquiring apical VSNs’ identity and AP-2ε expression specific to maturing (GAP43 positive) basal VSNs (Fig. 2C). As previously known [14], the expression of basal specific gene AP-2ε appears at later stages of maturation compared to apical specific Meis2 transcription factor. Following the indication of a previous study [11] that Bcl11b is a key gene in controlling cell fate choice of apical and basal VSNs, we looked at the spatial expression pattern of Bcl11b in Seurat object 3 (Fig. 2D). Overall, initial Bcl11b expression was seen after Ascl1 expression ended and after Neurog1/Neurod1 neuronal precursor stage started. On the basal VSN trajectory, Bcl11b expression was detected as a continuum from the dichotomy throughout the immature VSNs’ stage, whereas on the apical VSNs’ trajectory, Bcl11b mRNA expression was not detected until early stages of maturation (Arrows in Fig. 2D). Based on this observation, we manually clustered Bcl11b positive and negative cells specifically at the dichotomy (see boxed area in Fig. 2D and 2D’). Performing differential gene expression analysis between Bcl11b positive and negative cells specifically at the dichotomy (Fig. 2D’), we found Notch1receptor as an enriched gene in the Bcl11b positive cluster, whereas Dll4, which is a Notch ligand, is enriched in the Bcl11b negative cluster (Fig. 2D’’).

Moreover, we also identified Hes5, Hey1, Nrarp, and Ccnd1 enriched in the Bcl11b positive cluster (Fig. 2D’’), which along with Bcl11b are known to be downstream Notch signaling targets [31-34]. Spatial colocalization plots of Notch1 and Dll4 highlighted their mutually exclusive expression in cells before the neuronal dichotomy was established (Fig. 2E). Even though Notch mRNA could be detected in both apical and basal maturing VSNs after the dichotomy (Fig. 2E), Notch downstream targets like Hey1, Hes5, Nrarp and Ccnd1 appeared to be expressed only in Notch1+ but not Dll4+ cells (Fig. 2F) before the VSN bifurcation and in the early stage basal VSN trajectory (Fig. 2F). In addition, Dll4 ligand expression was found along with Meis2, which is an apical specific VSN marker, whereas downstream Notch signaling target Hey1 was found to be expressed along the basal VSN trajectory before Ap2ε (see colocalization feature plots in Fig. 2G, H). These gene expression plots suggest that Notch signaling is selectively activated in the cells acquiring basal identity during the establishment of the apical vs basal dichotomy but not after. Notably, many cells that are negative for Notch downstream target genes at the dichotomy are positive for Numb expression, suggestive of a post-translational Notch degradation after the dichotomy (Supplementary Fig. 2) [35, 36].

To further validate the biological relevance of differential Notch-Delta expression in the developing VSNs, we performed immunofluorescence staining against Dll4, Notch1, and Neurod1 on vomeronasal tissue at postnatal day 1 (P1). Double immunofluorescence of Dll4 and Notch1 confirmed the presence of immunodetectable Notch and Delta limited to Neurod1 positive precursors in the VNO (Fig. 3A, B, C). This analysis also highlighted Dll4+ and Notch1+ cells lying close to each other (Fig. 3A’’). These data suggest that differential Delta-Notch expression and activation of Notch signaling, in newly formed VSNs’ precursors, could play a role in establishing a binary differentiation of VSNs.

**Figure 3:**
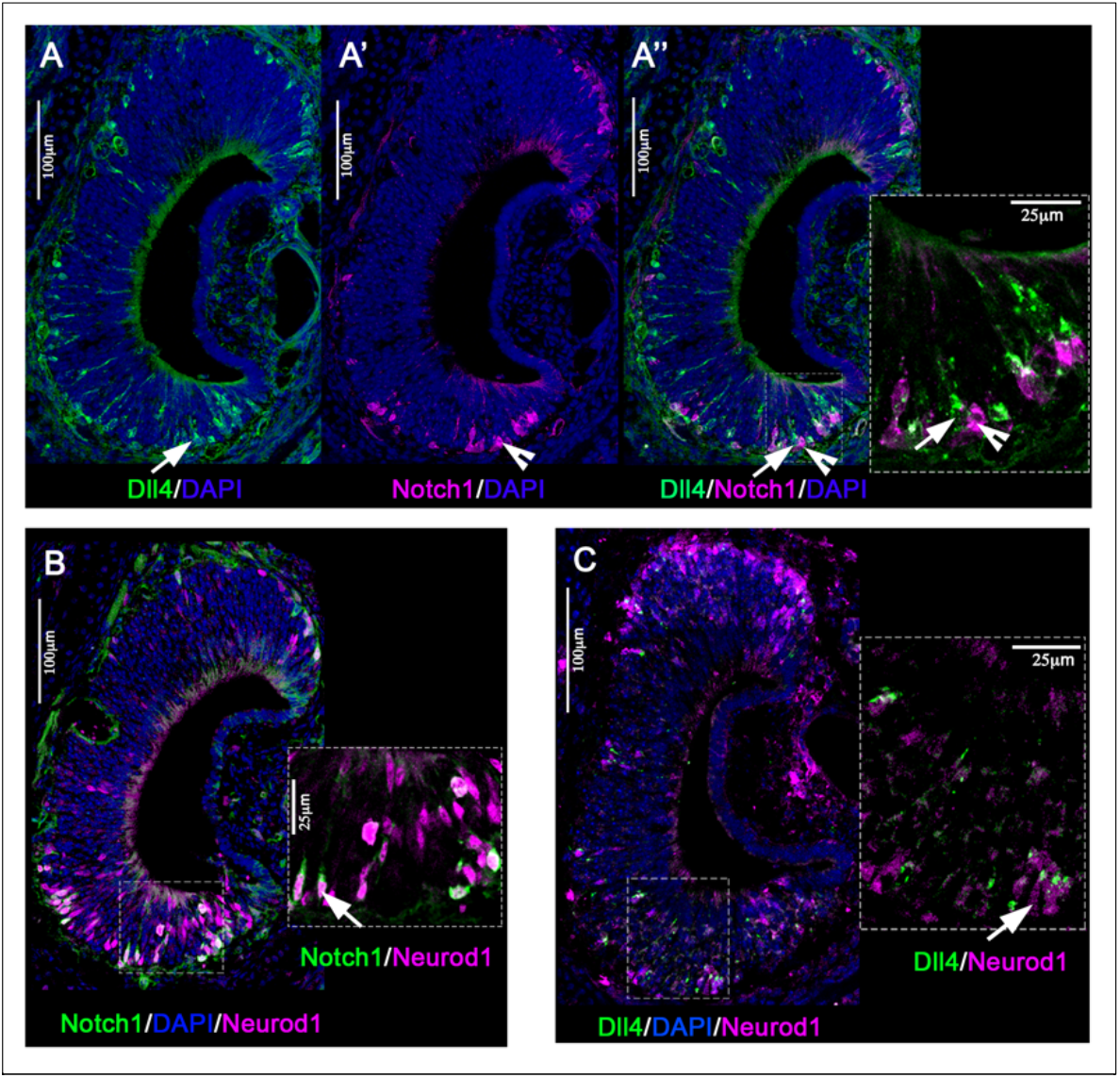
Notch1 and Dll4 immunoreactivity in Neurod+ cells. **A)** Double immunofluorescence of Dll4 in green and Notch1 in magenta shows expression of Notch ligand and receptor in marginal zones and basal regions of the VNO at postnatal day1. A’’ inset shows Dll4 (arrow) and Notch1+ (arrowhead) cells lying close to each other. **B)** Double immunofluorescence of Notch1 in green and Neurod1 in magenta shows (arrow) that Notch1 expression is mostly seen in Neurod1 stage. **C)** Double immunofluorescence of Dll4 in green and Neurod1 in magenta shows (arrow) that Dll4 ligand expression is mostly seen in Neurod1 stage.

### Conditional lineage tracing confirms the formation of Notch1+ and Dll4+ cells from Ascl1+ progenitors

Temporally controlled genetic lineage tracing of stem cell/progenitors can be used to follow proliferation and differentiation dynamics [37]. We performed conditional lineage tracing of Ascl1Cre^ERT2^/R26tdTom pups at P1 stage and collected VNOs at 1 day (1dpi), 3 days (3dpi) and 7 days post injection (7 dpi) (Fig. 4A). Further immunostainings were done to answer 2 main questions: 1) How long does it take for Ascl1 progenitors to become post mitotic? and 2) Do Ascl1+ progenitors give rise to both Dll4+ and Notch1+ VSNs’ precursors?

**Figure 4:**
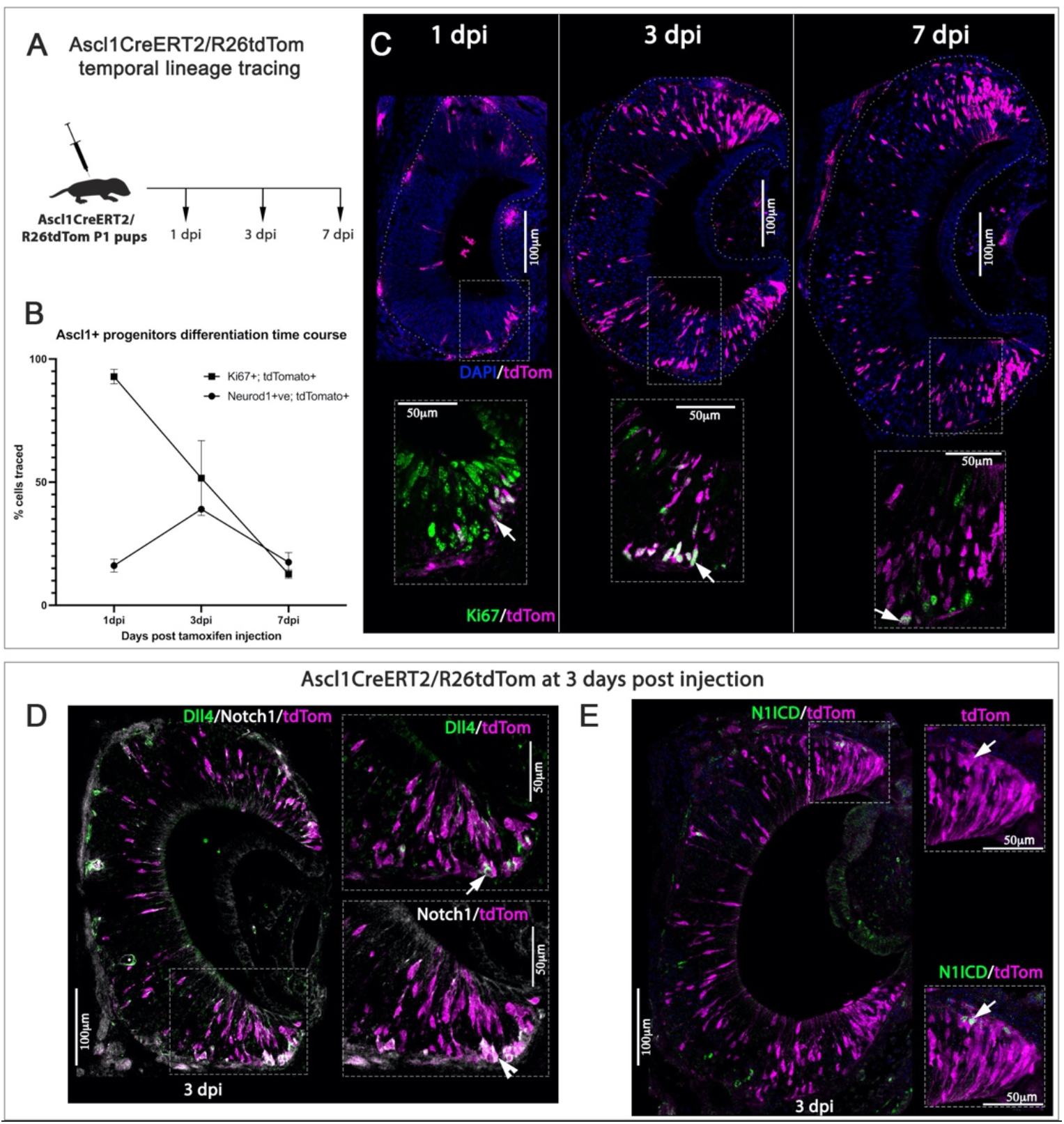
Ascl1 lineage tracing reveals that both Dll4 and Notch1+ cells are progeny of Ascl1+ cells. **A)** Schematic of experimental design for Ascl1Cre^ERT2^/R26tdTom lineage tracings. Tamoxifen injected to pups at P1 stage and perfused at 1 day post injection(1dpi), 3 days post injection (3dpi) and 7 days post injection (7dpi) **B)** Plot showing differentiation time course of Ascl1+ neuronal progenitors. % tdTom+ traced cells that are Ki67 positive and Neurod1 positive are quantified at 1dpi, 3dpi and 7dpi. **C)** Top panel shows number of tdTom+ traced cells increased from 1dpi to 3dpi and 7dpi. Bottom panel shows double immunofluorescence of proliferative marker Ki67 in green and tdTom in magenta. **D)** Triple immunofluorescence of Dll4, Notch1 and tdTom in Ascl1Cre^ERT2^ lineage traced pups at 3dpi stage. Inset shows both Dll4+ (arrow) and Notch+ (arrowhead) cells are colocalized with tdTom tracing. **E)** Double immunofluorescence of Notch Intracellular domain (NICD) in green and tdTom in magenta in Ascl1CreERT2 lineage traced pups at 3dpi stage. Inset shows tdTom+ cells colocalized with NICD (arrow) staining.

After a single tamoxifen injection, we observed that the number of Ascl1Cre^ERT2^ traced cells increased steadily from 30±4 cells at 1 dpi to 142±25 cells at 3dpi and 225±58 cells at 7 dpi as expected (n=3 at each stage and values in ± SD). However, the percentage of proliferative traced cells (Ki67+/tdTom) decreased from 93±2.9% at 1 dpi to 12±1.7% at 7 dpi (n=3 at each stage and values in ± SD) (Fig. 4B). These data suggest that Ascl1+ transit amplifying cells [38] of the VNO go through multiple rounds of cell division before becoming postmitotic neurons.

Analyzing the expression of Neurod1, which is a marker of cells undergoing differentiation (Fig. 1B’, 2B), we observed that Ascl1 traced cells, positive for Neurod1 immunoreactivity, transiently increased from 16±2.6% at 1dpi to 39±0.6% at 3 dpi and then decreased to 17±3.9% at 7dpi as they start to mature (n=3 at each stage and values in ± SD) (Fig. 4B). These results indicate that 1 week after temporally controlled Ascl1 lineage tracing, most of the cells become post mitotic and committed towards neuronal differentiation.

To further confirm the suitability of the experimental paradigm, we quantified the number of Ascl1Cre^ERT2^ traced cells that entered either the apical Meis2+ or basal Meis2-(See Fig. 2C) differentiation programs. Meis2/tdTom quantifications at 7dpi indicated that 51% of the traced cells entered the apical VSN differentiation program while the rest of the Meis2-traced accessed the basal VSN lineage (Fig. 5B, D). Overall, this experiment indicates that 7 days after Ascl1Cre^ERT2^ recombination is sufficient time interval to analyze the phenotypes emerging after genetic manipulations of vomeronasal progenitors.

**Figure 5:**
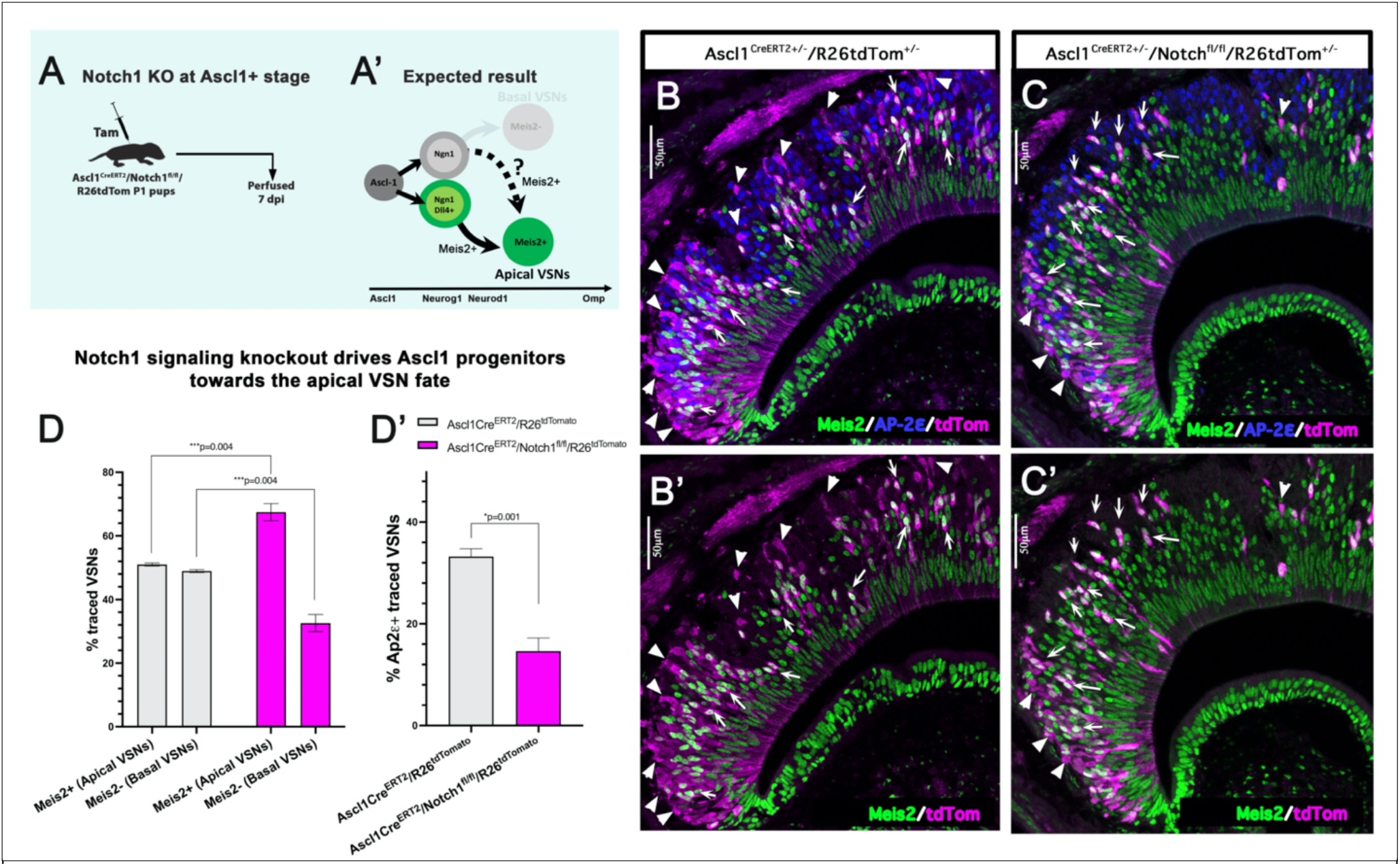
Notch1 LOF directs VSN differentiation to apical fate. **A)** Cartoon summarizing experimental design of Notch1 receptor loss of function study. Ascl1Cre^ERT2^/Notch1^fl/fl^/R26tdTom pups are injected with tamoxifen at P1 stage and perfused at 7dpi. **A’)** Cartoon of expected result showing Notch1 receptor knockout at Ascl1 stage may drive the progenitors towards apical VSN fate. **B, C)** Triple immunofluorescence of Meis2, Ap2ε and tdTom in control (B) and Notch1 KO (C) at 7dpi. **B’, C’)** Corresponding images of control and KO mice with only Meis2 and tdTom. Arrows highlight traced neurons that are Meis2 positive apical VSNs and arrow heads highlight traced neurons that are Meis2 negative basal VSNs. **D)** Bar graphs showing percentage of traced VSNs that are Meis2+ apical VSNs and Meis2-basal VSNs in control (gray) and Notch1 KOs (Magenta). **D’)** Bar graphs showing percentage of traced VSNs that express basal maturation marker Ap2ε in control and Notch1 KOs.

In addition to this, we also performed Dll4/Notch1/tdTom triple immunofluorescence staining for Ascl1Cre^ERT2^/R26tdTom tracing at 3dpi (Fig. 4D). As expected, we saw Dll4/tdTom and Notch1/tdTom double positive cells mostly in the marginal zones and basal zones of the vomeronasal sensory epithelium, where most of the postnatal neurogenesis takes place. Moreover, staining against cleaved Notch Intracellular Domain (NICD) confirmed Ascl1+ traced progeny undergoing active Notch signaling at 3 dpi (Fig. 4E). In summary, these results along with the Neurod1 immunofluorescence staining (Fig. 3B, C; Fig 4B) suggest that Ascl1+ progenitors divide further to give rise to Neurod1+ precursors that transiently express either Dll4 ligand or Notch1 receptor that can undergo active Notch signaling.

### In the absence of active Notch signaling maturing VSNs default to the apical cell fate

To test the role of Notch signaling in establishing the differentiation of VSNs, we decided to conditionally knock out Notch1 receptor from Ascl1+ progenitor stage onwards. We chose Ascl1Cre^ERT2^ driver, as both Notch1 and Dll4 expression start immediately after Ascl1 stage and around the Neurog1/Neurod1 stage (see Fig. 2B, 2E). We induced Cre recombination at P1 in both Ascl1Cre^ERT2^/R26tdTom^+/-^ controls and Ascl1Cre^ERT2^/Notch1^fl/fl^/R26tdTom^+/-^pups and analyzed at 7 days post injection. In control pups,1 week after lineage tracing, 51% of the traced cells were found positive for the apical marker Meis2+, while 49% cells were Meis2-(Fig. 5B, D). Analyzing the Ascl1Cre^ERT2^/Notch1 conditional KOs at 1 week after induction, we observed that Meis2+ apical population significantly increased from 51% in control to 67% in the conditional KO, whereas Meis2-population decreased proportionally. A deviation from the basal differentiation program was further confirmed with Ap2ε immunostaining which showed around 50% reduction in the maturing basal VSNs 1 week after conditional mutagenesis (Fig. 5D’). These results suggest that active Notch signaling, via Notch1 receptor, is essential to trigger the activation of the basal differentiation program.

### Conditional induction of NICD at Ascl1 progenitor stage pushed lineage towards sustentacular cell formation

Ablation of Notch signaling suggests that without Notch signaling VSN progenitors’ default towards the apical VSN cell fate. To further test whether active Notch signaling is sufficient to initiate basal VSN genetic program, we decided to conditionally overexpress NICD at Ascl1 stage, using Ascl1Cre^ERT2^/R26NICD^+/-^ pups. By doing so, we aimed to test if activation of Notch signaling would redirect all the recombined cells towards the basal VSN phenotype. In the inducible experiment, we counted GFP positive cells to follow NICD recombined cells as these mice have Cre^ERT2^ activated NICD along with nuclear localized EGFP reporter [39]. The total number of GFP+ NICD recombined cells are less compared to control Ascl1Cre^ERT2^/R26tdTom tracing which can be due to low efficiency in R26NICD recombination as previously reported [40].

We induced Cre recombination at P1 in both control Ascl1Cre^ERT2^/R26tdTom^+/-^ and Notch inducible Ascl1Cre^ERT2^/R26NICD^+/-^pups and analyzed for the phenotype at 7 days post injection. In the control mice, tdTom/Meis2 immunofluorescence was performed to quantify non-neuronal sustentacular cells and, apical and basal VSNs. While Meis2 is expressed in both apical VSNs and sustentacular cells, the morphology of the tdTom traced cell is used to distinguish between sustentacular cell and apical VSN. The analysis showed that at 7 dpi in control tracing, 99.3% of traced cells became VSNs out of which, ∼50 % expressed the apical marker Meis2 and less than 1% of the traced cells were non-neuronal Sustentacular cells (Fig. 6B, F).

However, in Ascl1Cre^ERT2^/R26NICD^+/-^ pups, we found, to our surprise, conditional NICD overexpression in Ascl1+ progenitors mostly induced the formation of non-neuronal Sustentacular cells. We performed GFP/Meis2/HUCD triple immunofluorescence to distinguish among sustentacular cells and, apical and basal VSNs in the NICD inducible pups. As HUCD is only expressed in neuronal population [41], we considered Meis2+/HUCD-as non-neuronal sustentacular cells, Meis2+/HUCD+ as apical VSNs and Meis2-/HUCD+ as basal VSNs. The analysis in the inducible mice showed that 66.6% of the NICD: EGFP population became sustentacular and only 32.3% of the recombined cells were found to be VSNs (Fig. 6C, F). Within the recombined neuronal population, nearly all of them were negative for the apical marker Meis2 (Fig. 6F’). These data suggest that sustained activation of NICD in Ascl1 progenitor mostly deviates the neurogenic progenitors towards Sustentacular cells differentiation. However, the neurons that formed from NICD+ Ascl1 progenitors resulted to be Meis2 negative, therefore not committed to apical fate.

**Figure 6:**
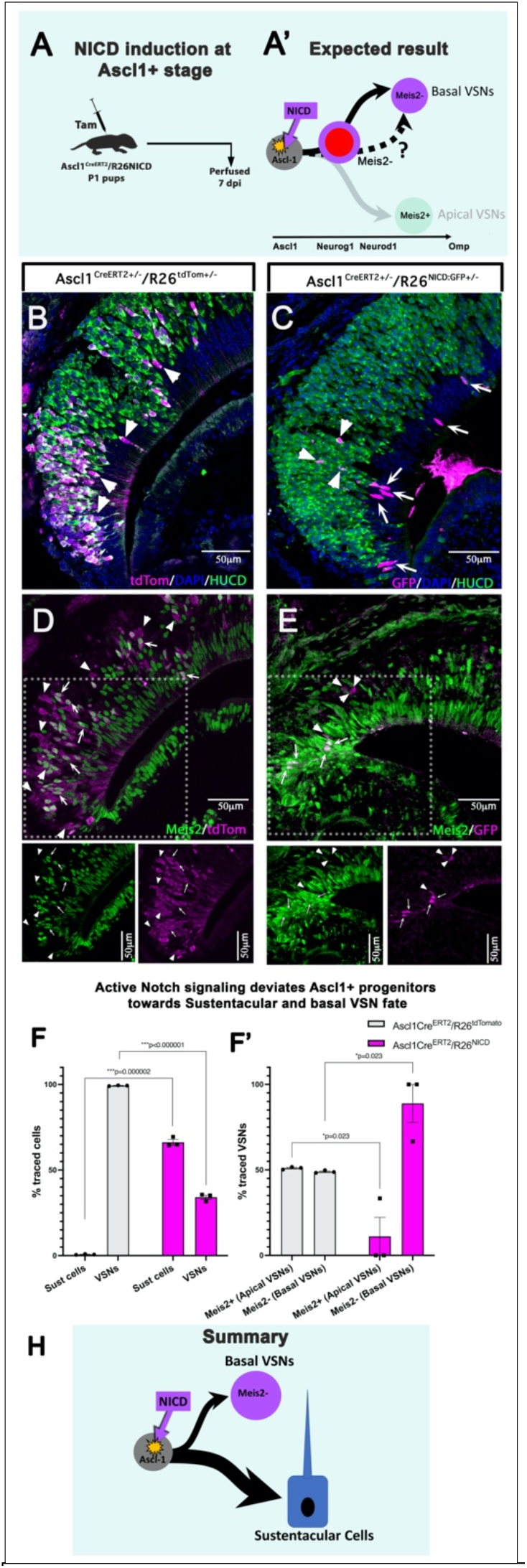
Ectopic expression of NICD at Ascl1 stage diverts the progenitors towards Sustentacular cell fate. **A)** Schematic showing experimental design of NICD gain of function study. Ascl1Cre^ERT2^/R26NICD pups are injected with tamoxifen at P1 stage and perfused at 7dpi. **A’)** Cartoon of expected result showing NICD overexpression at Ascl1 stage may drive the progenitors towards basal VSN fate. **B, C)** Double immunofluorescence of tdTom / HUCD in control and GFP/HUCD in NICD inducible mice at 7dpi. Arrows highlight NICD+/HUCD-sustentacular cells, whereas arrowheads highlight NICD+/HUCD+ VSNs. **D, E)** Double immunofluorescence of tdTom/Meis2 in control and GFP/Meis2 in NICD inducible mice at 7dpi. Arrows highlight traced Meis2+ apical VSNs in control mice and Meis2+ sustentacular cells in inducible mice. Arrow heads highlight Meis2-basal VSNs in both control and inducible mice. **F)** Bar graph plot showing %traced tdTom+ cells or GFP+ cells that are sustentacular cells or VSNs in control and NICD mice respectively. **F’)** Bar graphs showing %traced tdTom+ cells or GFP+ cells that are Meis2+ apical VSNs or Meis2-basal VSNs in control and NICD mice respectively. **H)** Cartoon of summary results showing NICD overexpression at Ascl1 stage mostly differentiate progenitors into non-neuronal Sustentacular cells. Within the neuronal population most of the NICD+ cells deviate towards basal VSN cell fate.

### Conditional induction of NICD at Neurog1 precursors stage leads to reduction in Meis2+ apical VSN formation

Based on the intriguing results obtained at Ascl1 stage, we decided to test if later NICD activation, in committed (Neurog1+) neuronal precursors, could give us a clearer readout of the role of NICD in controlling the apical-basal dichotomy. Moreover, the Neurog1/Neurod1 stage is coincident to the stage when Notch signaling is more active and the dichotomy appears to be established (Fig 2 B, D, E, G).

We induced Cre recombination at P1 in both control Neurog1Cre^ERT2^/R26tdTom^+/-^ and Notch inducible Neurog1Cre^ERT2^/R26NICD^+/-^ pups and analyzed 7 days post injection. We counted tdTomato+ cells and NICD: EGFP+ cells to follow the recombined cells in control and inducible R26NICD mice respectively. In control tracing experiment, Meis2/tdTom analysis at 7dpi showed around 52% of the cells expressed apical VSN marker (Meis2+) while the rest of the 48% population acquired basal (Meis2-) VSN fate (Fig. 7 B, D). Notably, within the Meis2-cells that enter the basal program, 96.57% expressed the transcription factor AP-2ε suggesting that they arrive to the basal maturing stage.

**Figure 7:**
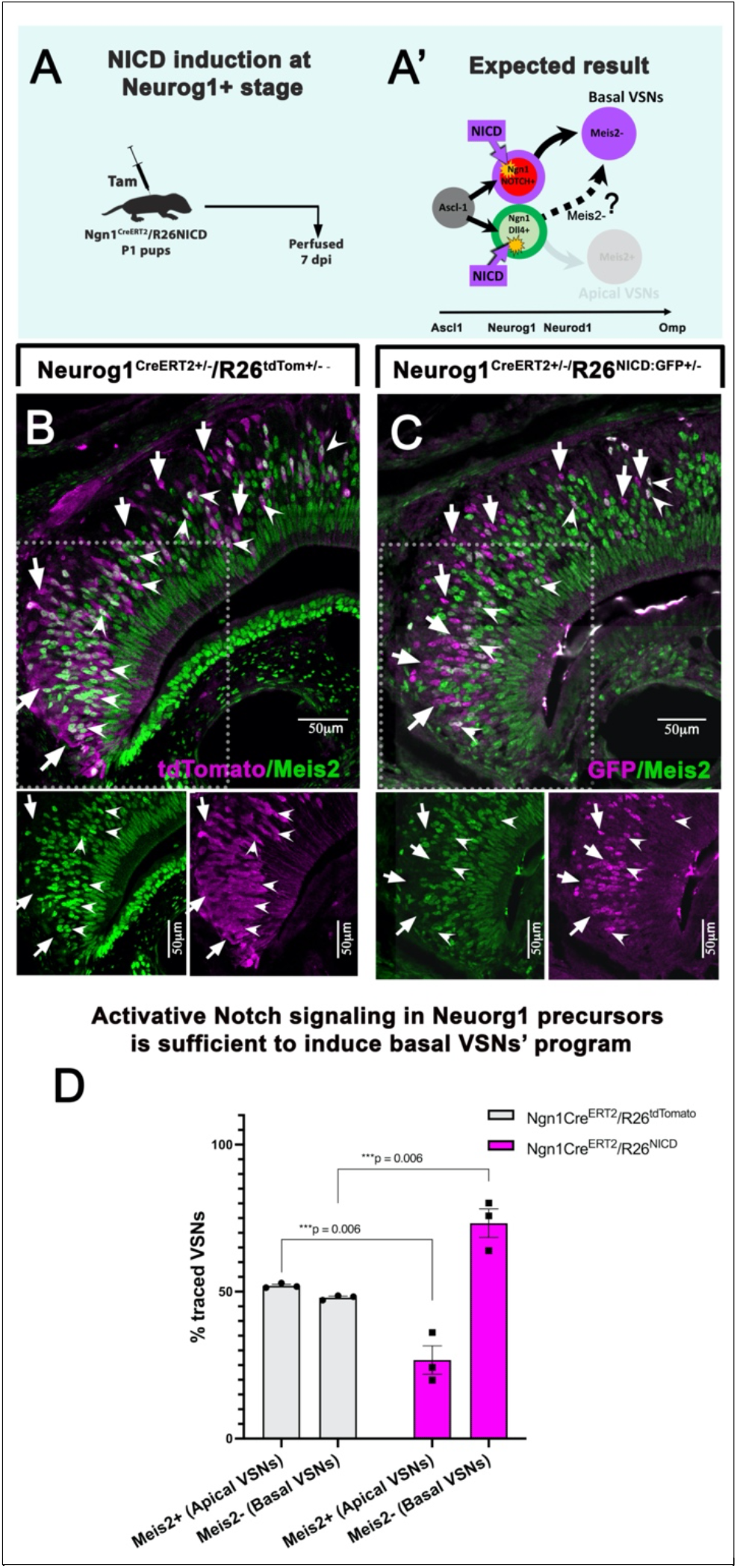
Ectopic expression of NICD at Neurog1 stage diverts the neuronal precursors towards basal VSN fate A) Schematic showing experimental design of NICD gain of function study. Neurog1Cre^ERT2^/R26NICD pups are injected with tamoxifen at P1 stage and perfused at 7dpi. **A’)** Cartoon of expected result showing NICD overexpression at Neurog1 stage may drive the progenitors towards basal VSN fate. **B, C)** Double immunofluorescence of tdTom/Meis2 in control and GFP/Meis2 in NICD inducible mice at 7dpi. Arrows highlight traced Meis2-basal VSNs and arrowheads highlight Meis2+ apical VSNs in both control and inducible mice. **D)** Bar graphs showing %traced tdTom+ cells or GFP+ cells that are Meis2+ apical VSNs or Meis2-basal VSNs in control and NICD mice respectively.

However, in the Neurog1Cre^ERT2^/R26NICD inducible mice, one week after constitutive NICD overexpression, we found that Meis2-putative basal VSN population significantly increased to ∼ 73% (Fig. 7 C, D). Interestingly, within this Meis2-population, only 56.4% expressed the basal maturation marker AP-2ε. These data suggest that induction of Notch signaling at Neurog1 stage is sufficient to deviate the newly formed VSNs’ away from the apical (Meis2+) differentiation program, however, prolonged NICD signaling might also interfere with normal basal VSN maturation.

## Discussion

Interaction of extrinsic and intrinsic regulators is important for neural stem cells/ progenitors to make cell fate decisions and give rise to specific neuronal cell types. The VNE of mice is composed of V1R+/Gαi2+ apical and V2R+/Gαo+ basal VSN sub types. Correct development and function of these two main types of vomeronasal neurons is important for social, sexual, maternal and predator avoidance behaviors of rodents [16-19]. However, Bcl11b is the only known intrinsic regulator till now shown to affect the cell fate decision in the VNO [11]. In the current study, for the first time we identified the role of Dll4-Notch1 signaling in the cell fate specification of apical and basal VSNs. In addition, research from T cell development previously reported Bcl11b as direct downstream Notch signaling target gene [42], thus relating our current study to the Bcl11b role in the VNO.

In this study, sc-RNA seq analysis of adult VNO at the neuronal dichotomy identified Dll4 ligand and Notch1 receptor in the apical and basal committed neuronal precursors respectively. In addition, immunofluorescence data along with the conditional Ascl1Cre^ERT2^ lineage tracing also confirmed that both Notch1+ and Dll4+ cells are the progeny of Ascl1 cells and are expressed at Neurod1 stage, suggesting a potential role for Dll4-Notch1 signaling in establishing the VSN dichotomy. Previous studies also highlighted the expression of Notch1 and Dll4 in developing VNO at different ages [43, 44].

Notch is an evolutionarily conserved juxtracrine signaling pathway associated with inhibitory interactions/lateral inhibitions that can determine cell fates among two juxtaposed differentiating cells [45, 46]. Notch inhibitory ligand-receptor interactions rely on non-symmetric expression of a Notch transmembrane receptor and a Notch ligand between neighboring cells. Notch activation in a cell prevents it from assuming the same cell fate as the neighboring cell expressing the Notch ligand. Notch paralogs (Notch1, Notch2, Notch3, Notch4), three Delta-like ligands (Dll1, Dll3, Dll4) and two Jagged-like ligands (Jagged1, Jagged2) have been found in mammals [47]. In this study, we performed LOF and GOF experiments at Ascl1+ neuronal progenitor stage and Neurog1+ neuronal precursor stage to probe the necessity and sufficiency of Notch signaling in establishing neuronal diversity in the VNO. In the LOF strategy, Notch1 receptor knockout from Ascl1+ neuronal progenitor stage onwards caused most of the neurons to enter the apical (Meis2+/V1R/Gai2) differentiation program. This suggests that neuronal progenitors are by default fated towards apical VSN type if not instructed otherwise by Notch mediated transcriptional regulation. Vomeronasal precursors with active Notch signaling express Bcl11b [11] and then AP-2ε at a later maturation stage [14]. From our single cell seq analysis, we found that Ascl1+ progenitors don’t express Notch1 receptor. In line with this, Notch1 ablation at Ascl1 stage does not affect the neurogenic process, but the cell fate choice of VSNs.

Notch pathway has been extensively shown to be important for promoting gliogenesis and for maintaining glial function in rodents [48-51]. Moreover, ectopic activation of Notch signaling has also been previously shown to be able to alter cell fate [52]. Ectopic expression experiments of transcription factors at different maturation stages of neurons have revealed that the dynamic changes in the chromatin landscape play a key role in restricting the phenotypic plasticity of the cells [53]. In line with this, our conditional ectopic NICD GOF experiments in the VNO gave distinct phenotypes based on the timing of Cre recombination. In fact, NICD activation in Ascl1+ neural committed progenitors, which in control conditions mostly give rise to VSNs (see Fig. 6F), appears to be sufficient to induce their differentiation to Sustentacular cells. This non-neuronal phenotype may likely result from NICD-mediated negative regulation of pro-neurogenic transcription factors [54, 55] and by the activation of the alternative Sustentacular differentiation program. Notably, we already know that NICD expression in multipotent olfactory horizontal basal cells (HBCs) during regeneration is sufficient to induce Sustentacular cell differentiation and that Notch signaling is required for homeostasis of the Sustentacular cells [48, 56, 57]. Moreover, sc-RNA seq data also identified the expression of downstream Notch targets in the VNO Sustentacular population suggesting a role of Notch signaling in their formation (Supplementary Fig. 3). Notably, Ascl1+ progenitors have more restricted potency compared to the HBCs [58]. However, the phenotypic change of vomeronasal Ascl1+ progenitors in response to NICD implies that, at this stage, the chromatin landscape is still plastic enough to be rearranged by active Notch signaling[53, 59].

On the other hand, NICD activation at Neurog1 stage, did not convert neuronal cells to Sustentacular cells, but increased the proportion of Meis2-basal VSNs. This suggests that at Neurog1 stage, the VSNs’ precursors reach a level of commitment at which the chromatin landscape is no longer permissive [53] for Notch-mediated reprograming from neurons to glia/Sustentacular cells. However, at this stage Notch signaling is sufficient to inhibit the progression of the Meis2+ apical VSN program and to divert the neuronal precursors towards the basal VSN fate. One interesting aspect with the ectopic GOF experiments is the relation of NICD with Meis2 expression. Meis2 is expressed in both apical VSN lineage and non-neuronal Sustentacular cells. Ectopic NICD expression at Ascl1 stage not only converted them into Sustentacular cells, but also maintained their Meis2 expression, whereas NICD overexpression at Neurog1 stage repressed apical VSN specific Meis2 expression diverting them into basal lineage. This suggests that NICD complex follows different downstream neuronal and sustentacular differentiation pathways depending on the developmental stage of the cell.

From our sc-RNAseq data (see Fig. 2E, 2G) and Ascl1Cre^ERT2^/R26tdTom control tracing (see Fig. 4E) we know that, in physiological conditions, active Notch signaling is temporally restricted to Notch1+ cells when the apical-basal dichotomy is established. After Neurog1Cre driven NICD expression, we found that even though more VSNs deviate from the apical program, a smaller percentage of Meis2-cells (basal committed) were able to express the basal maturation marker Ap2ε. This reduction in the Ap2ε expression may likely result from the non-physiological timing/levels of NICD activation in the Neurog1^CreERT2^/R26NICD mice.

Notch signaling role in cell fate specification is conserved in many neuronal and non-neuronal systems [60-63]. Further, research into Notch1-Dll4 signaling literature in retina and spinal cord pointed out Foxn4, a forkhead transcription factor to have a role in inducing Notch1-Dll4 mediated signaling [63-65]. In retina, Foxn4 is important in diverting the retinal progenitor cell fate towards amacrine cells and horizontal cells, whereas in spinal cord, Foxn4 is expressed in p2 progenitors that specifically give rise to V2a and V2b interneurons. In both systems, it has been shown that Foxn4 controls the expression of Dll4 and that Foxn4 mutants have aberrant cell fate phenotypes [63, 64]. Our sc-RNA seq data revealed Foxn4 mRNA expression partially overlapping to that of Dll4. We further corroborated co-expression via Dll4/Foxn4 double immunofluorescence staining at E14 stage (Supplementary Fig. 4). The identification of Dll4 and Foxn4 double positive cells suggested to us to test whether Foxn4 plays a role in apical vs basal VSN differentiation via Dll4 ligand expression. However, Foxn4 null mutants still showed, as the wild type, detectable Dll4 expression at E14 stage, moreover, no obvious changes in the apical-basal VSN ratio was detected (Supplementary Fig. 4). These data suggest, that in the VNO, Foxn4 plays a dispensable role in inducing Dll4 expression.

Among Notch modulators, mammalian Numb and Numb-like proteins have been shown to repress Notch signaling by controlling NICD endosomal trafficking and degradation [66]. By taking a closer look at downstream Notch signaling targets like Hey1, Hes5 and, CyclinD1 vs Numb in sc-RNA seq data, we confirm that Numb expression negatively correlates with Notch-mediated signaling in the VNO (Supplementary Fig. 2). Notably these molecules play important roles in modulating the efficiency of Notch signaling but not Notch or delta expression.

By analyzing our single cell data, we observed that Notch and Delta mRNA appeared to be non-symmetrically distributed and mutually exclusive among cells even before the apical-basal VSNs’ dichotomy is established (Fig. 2E). These data suggest that non-symmetric inheritance of RNA binding proteins or other post-transcriptional modulators [67-69] define the predetermination of the Notch and Delta neuronal precursors. In support of this idea, we observed that KH domain containing RNA binding protein (RBP) Oocyte expressed protein (Ooep), was significantly enriched in the Dll4+/Bcl11b-cluster but not Notch1/Bcl11b+ cluster at the VSN dichotomy (Fig. 2). Ooep, also named as Floped/Moep19 is a maternal effect gene that is expressed in oocyte and regulates embryonic cell division via F actin cytoskeleton [70-73]. Moreover, in our data, even after the dichotomy, Ooep expression was found to persist only in the apical VSN branch along with Meis2 which is one of the early apical VSN marker (Supplementary Fig. 5). As RBPs can also control cell fate choice by post-transcriptionally modulating mRNA and miRNA dynamics [74-76], it would be interesting to further study whether Ooep can work as intrinsic regulator of the Notch and Delta asymmetry in neuronal precursors.

Evolution of new sensory neuronal types in animals can have an important role in determining their social and environmental fitness by expanding their ability to detect, compute, and respond to new stimuli. Both V1R and V2R are functionally, and evolutionarily unrelated super families of receptors and previous studies highlighted the diversity of V1R and V2R positive VSN population across many vertebrate species [77, 78]. In most of the mammals e.g., horse, goat, musk shrew, common marmoset, dog, and cow, few to none V2R+ genes appear to be expressed. Notably rodents and opossum have strikingly expanded repertoire of functional V2R genes, moreover in these animals, specialized VSNs segregate into V1R+/Gαi2+ apical and V2R+/Gαo+ basal VSNs [10, 79]. As the current study has revealed the role of Notch signaling in the VSN cell fate specification, it would be interesting to further study the evolutionary development of mechanisms controlling Notch1-Dll4 expression across different vertebrate species and see how these mechanisms influence the neuronal composition of the chemosensory epithelia across species.

## Materials and Methods

### Mouse lines

We purchased Ascl1Cre^ERT2^ (*Ascl1*^*tm1*.*1(Cre/ERT2)Jejo*^/J Stock No: 012882), Neurog1Cre^ERT2^ (B6;129P-Tg(Neurog1-cre/ERT2)1Good/J Stock No: 008529), Notch1^fl/fl^(B6.129X1-*Notch1*^*tm2Rko*^/GridJ Stock No: 007181), R26NICD(STOCK *Gt(ROSA)26Sor*^*tm1(Notch1)Dam*^/J Stock No: 008159), R26tdTom (B6.Cg-Gt(ROSA)26Sortm9^(CAG-tdTomato)Hze^/J, JAX stock #007909) mouse lines from Jackson Lab. The *Foxn4*^*+/lacZ*^ mouse line was generated previously [80]. Genotyping was conducted following the suggested primers and protocols from JAX. Mice of either sex were used for immunohistochemistry and immunofluorescence experiments. All experiments involving mice were approved by the University at Albany Institutional Animal Care and Use Committee (IACUC).

### Single-Cell RNA Sequencing

The vomeronasal organs of P60 C57BL/6J male mice were isolated and dissociated into single-cell suspension using neural isolation enzyme/papain (NIE/Papain in Neurobasal Medium with 0.5mg/mL Collagenase A, 1.5mM L-cysteine and 100U/mL DNAse I) incubated at 37°C. The dissociated cells were then washed with HBSS and reconstituted in cell freezing medium (90% FBS, 10% DMSO). Cells were frozen from room temperature to -80°C at a -1°C/min freeze rate. The single cell suspension was sent to SingulOmics for high-throughput single-cell gene expression profiling using the 10x Genomics Chromium Platform. Data was analyzed using Seurat 3.15.

### Quality control and cell clustering

Quality control (QC), and clustering and downstream analysis was performed using Seurat [3.15] package in R. Basic filtering was carried out where all genes expressed in ≥3 cells and all cells with at least 200 detected genes were included. QC was based on number of genes and percent mitochondrial genes, where all cells that expressed > 9000 genes and > 5% mitochondrial genes were not included in the analysis. After filtering, 10,582 cells were included for the clustering and analysis. Top 2000 highly variable genes across the population were selected to perform PCA and the first 35 principal components were used for cells clustering, which was then visualized using uniform manifold approximation projection (UMAP). Stem cells, neuronal progenitors, precursors and immature neuronal cell types were identified based on the expression of known genes. These cell types were specifically chosen to subset and top 2000 highly variable genes and top 15 principal components were used to cluster and create new Seurat object2. Similarly, Seurat object3 was created by focusing on clusters only at VSN dichotomy and top 20 principal components were used to cluster and visualize. All downstream analysis that identified Notch1-Dll4 signaling were done on Seurat object3.

### Pseudotime analysis of cell population spatial organization

To further confirm the VSN dichotomy, we used Monocle3 to perform pseudotime analysis, where Ascl1 positive cells were chosen as root node. We used Seurat wrappers package to directly convert Seurat object2 into cell data set format.

### Tamoxifen Treatment

Tamoxifen (Sigma–Aldrich), CAS # 10540−29−1, was dissolved in Corn Oil at 20mg/ml concentration. For all tamoxifen inducible experiments, we injected tamoxifen once intraperitoneally at postnatal day 1 at a dose of 80mg/Kg body weight and perfused at indicated postnatal days.

### Tissue Preparation

Tissue collected were perfused with PBS followed by 3.7% formaldehyde in PBS. Noses were immersion fixed in 3.7% formaldehyde in PBS at 4°C for 1-2 hr depending on the age. All samples were cryoprotected in 30% sucrose in PBS overnight at 4°C then embedded in Tissue-Tek O.C.T. Compound (Sakura Finetek USA, Inc., Torrance CA) using dry ice, and stored at -80°C. Tissue was cryosectioned using a CM3050S Leica cryostat at 16μm for VNOs and collected on VWR Superfrost Plus Micro Slides (Radnor, PA) for immunostaining. All slides were stored at -80°C until ready for staining.

### Immunofluorescence

Citrate buffer (pH 6.0) antigen retrieval was performed [14], for all the antibodies indicated with asterisks (*). Primary antibodies and concentrations used in this study were, *Goat anti-AP-2ε (2ug/mL, sc-131393 X, Santa Cruz, Dallas, TX), Chicken anti-GFP (1:3000, ab13970, Abcam), Rabbit anti-GFP (1:1000, A-6455, Molecular Probes, Eugene, OR), *Rabbit anti-Ki67 (1:1000, D3B5, Cell signaling Tech), *Mouse anti-Ki67 (1:500, 9449, Cell signaling Tech), *Mouse anti-Meis2 (1:500, sc-515470, Santa Cruz), Rabbit anti-Meis2 (*with antigen retrieval 1:1000 and without antigen retrieval 1:500, ab73164, Abcam), *Mouse anti-NeuroD1 (1:100, sc-46684, Santa Cruz), *Goat anti-NeuroD1 (1:500, AF2746, R&D Systems), *Rabbit anti-Notch1 (1:50, D1E11, Cell Signaling Tech), Goat anti-Dll4 (*with antigen retrieval 1:50 and without antigen retrieval 1:25, AF1389, R&D Systems), *Rabbit anti-NICD activated (1:75, D3B8, Cell Signaling Tech), *Mouse anti-DsRd (1:500, TA180084, Origene), *Rabbit anti-DsRd (1:500, 600-401-379, Rockland), HuC/D 8 μg/ml (Molecular Probes), Rabbit anti-Foxn4 (1:50) [80].

For chromogen-based reactions, tissue was stained as previously described [14]. Staining was visualized with the Vectastain ABC Kit (Vector, Burlingame, CA) using diaminobenzidine (DAB); sections were counterstained with methyl green.

Species-appropriate secondary antibodies conjugated with either Alexa Fluor 488, Alexa Fluor 594, Alexa Fluor 568, Alexa Fluor 680 plus were used for immunofluorescence detection (Molecular Probes and Jackson ImmunoResearch Laboratories, Inc., Westgrove, PA). Sections were counterstained with 4’,6’-diamidino-2-phenylindole (DAPI) (1:3000; Sigma-Aldrich) and coverslips were mounted with FluoroGel (Electron Microscopy Services, Hatfield, PA). Confocal microscopy pictures were taken on a Zeiss LSM 710 microscope. Epifluorescence pictures were taken on a Leica DM4000 B LED fluorescence microscope equipped with a Leica DFC310 FX camera. Images were further analyzed using FIJI/ImageJ software.

### Experimental design, quantification, and statistical analyses of microscopy data

All data were collected from mice kept under similar housing conditions, in transparent cages on a normal 12 hr. light/dark cycle. Tissue collected from either males or females in the same genotype/treatment group were analyzed together unless otherwise stated; ages analyzed are indicated in text and figures. Measurements of VNE and cell counts were performed on confocal images of coronal serial sections immunostained for the indicated targets. Measurements and cell counts were done using ImageJ. The data are presented as mean ± SEM unless otherwise specified. Prism 9.0.1 was used for statistical analyses, including calculation of mean values, and SEM. Two-tailed, unpaired t-test were used for all statistical analyses, and calculated p-values <0.05 were considered statistically significant. Sample sizes and p-values are indicated as single points in each graph and/or in figure legends.

## Author contributions

PEF conceptualized and designed the experiments. RRK designed, performed the experiments, and analyzed the data. MX designed experiments. SL performed the experiments and analyzed the data. EZT, SMH analyzed the data. JML performed the experiments. RRK and PEF wrote the manuscript and designed illustrations.

## Funding

This publication was supported by the Eunice Kennedy Shriver National Institute of Child Health and Human Development of the National Institutes of Health under the Awards R15-HD096411 (P.E.F) and R01-HD097331/HD/NICHD (P.E.F), and by the National Institute of Deafness and Other Communication Disorders of the National Institutes of Health under the Award R01-DC017149 (P.E.F).

Mengqing Xiang’s lab contribution was supported by the National Natural Science Foundation of China (81970794), Natural Science Foundation of Guangdong Province of China (2020A1515010328)and the Fundamental Research Funds of the State Key Laboratory of Ophthalmology, Sun Yat-sen University.

## Supplementary figures

**Supplementary figure 1:**
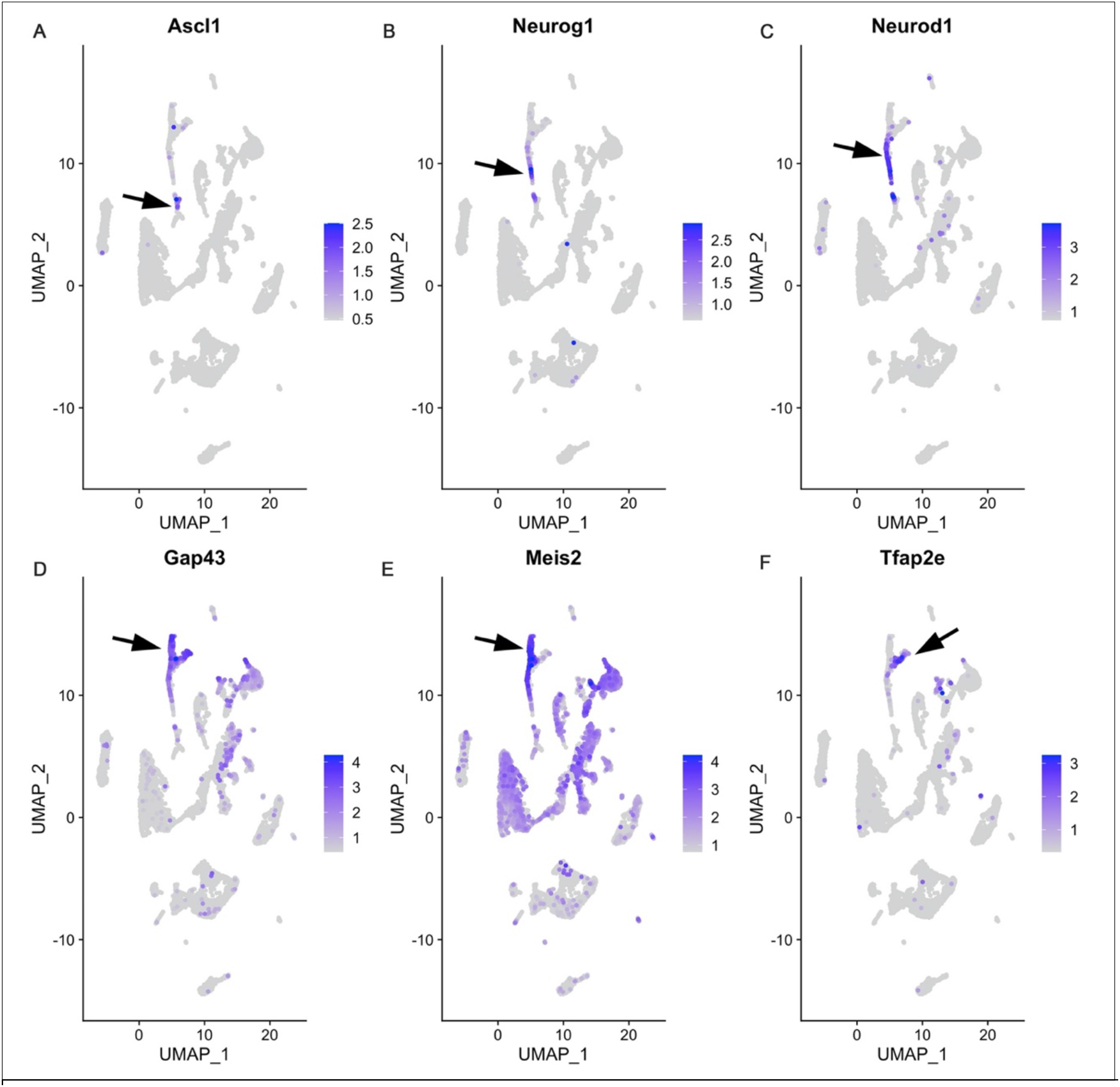
sc-RNA seq analysis identifies apical and basal VSN dichotomy. **A-F)** Feature plots of Ascl1, Neurog1, Neurod1, Gap43, Meis2 and Ap2ε genes that are specific to neuronal progenitors, precursors and immature apical and basal VSNs identify VSN dichotomy. Specific expression of the gens is highlighted by arrow. Both Meis2 and Tfap2e expression highlights apical and basal VSN branches respectively.

**Supplementary figure 2:**
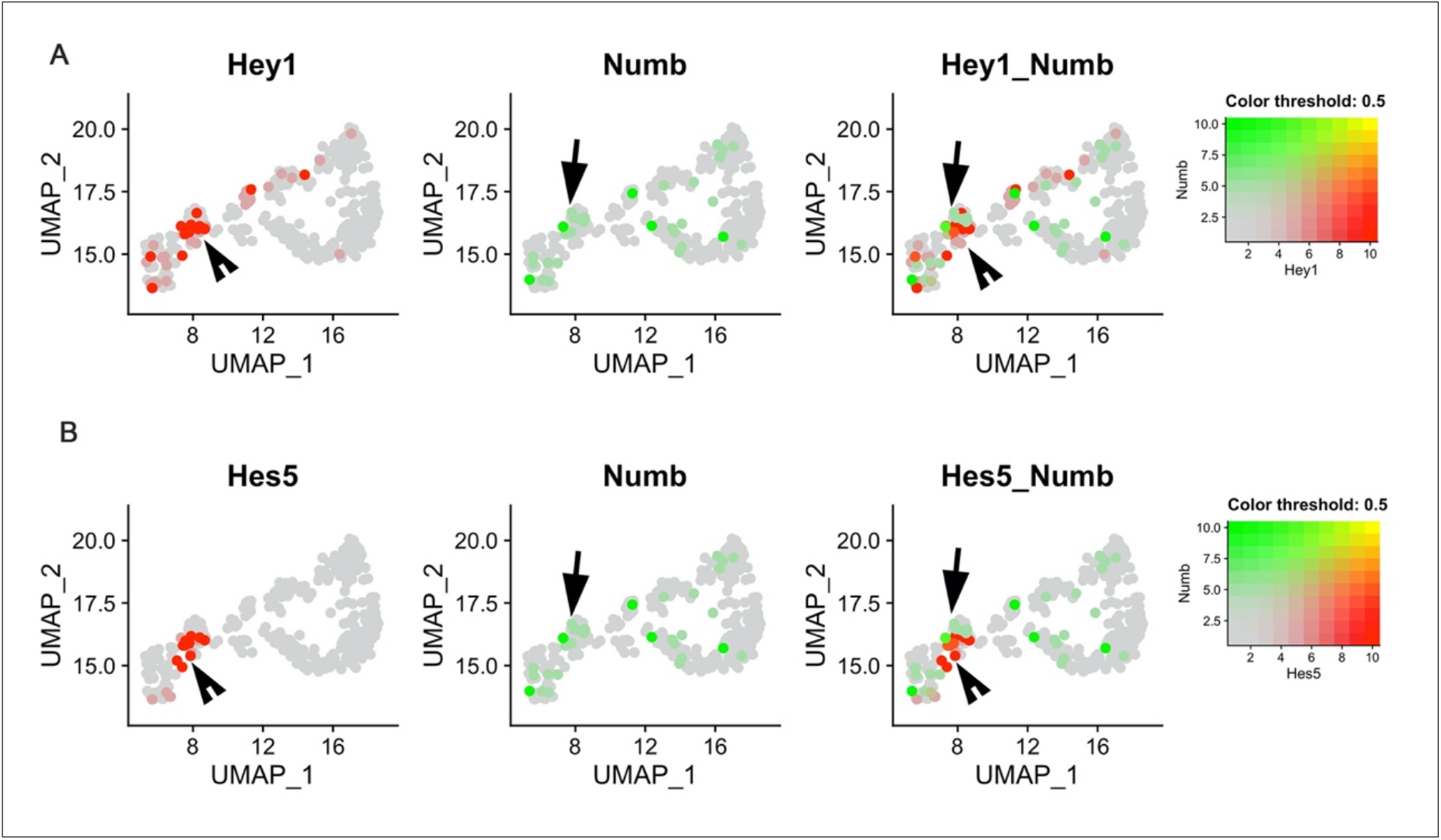
Expression of Numb and downstream Notch targets are mutually exclusive at the VSN dichotomy. **A, B)** Blended feature plots of Notch target genes Hey1 and, Hes5 Vs Numb shows that Notch target genes are not expressed in similar populations along with Numb at the VSN dichotomy.

**Supplementary figure 3:**
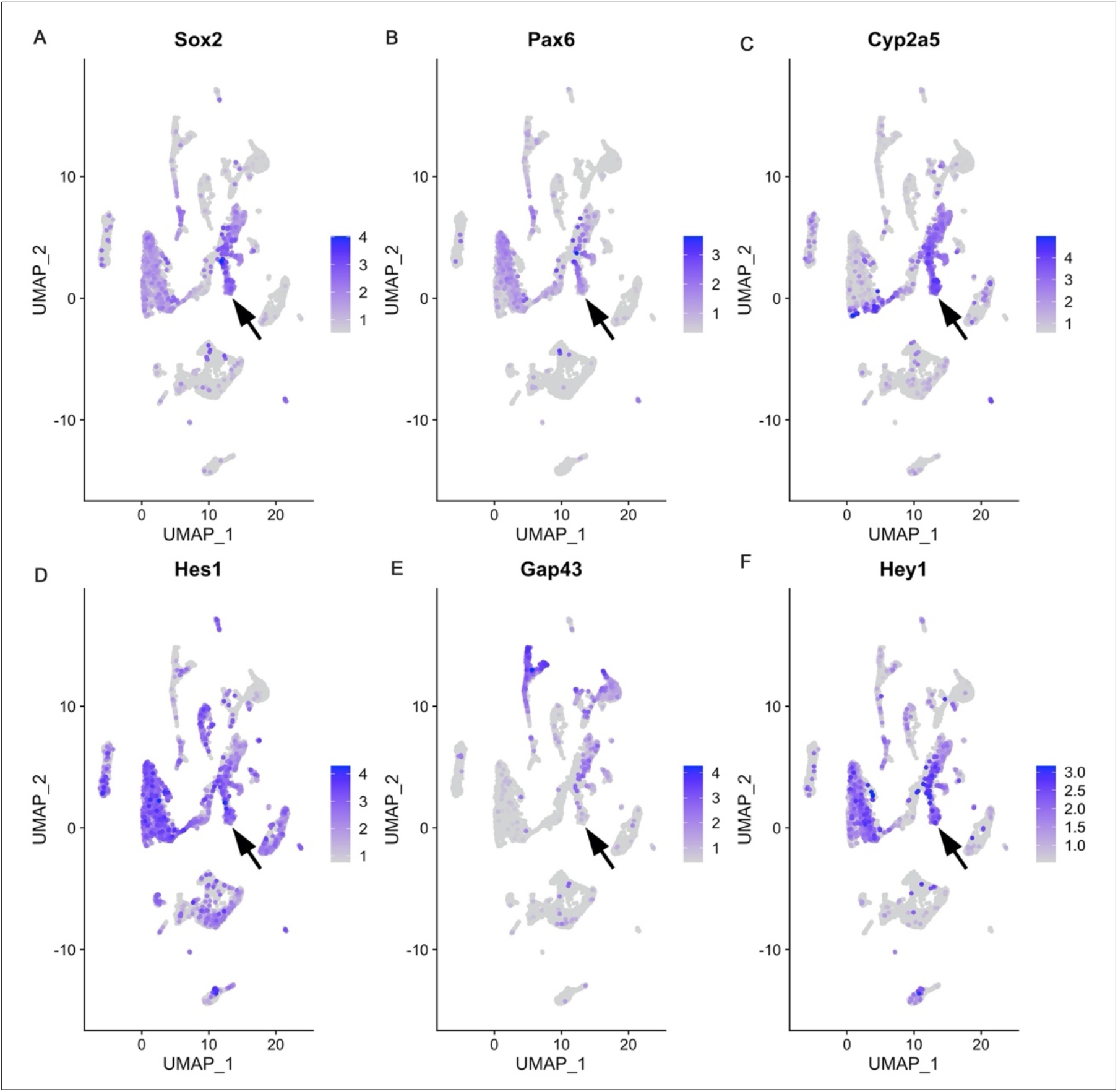
sc-RNA seq analysis reveals the expression of downstream Notch targets in Sustentacular cells. **A-D)** Feature plots of Sox2, Pax6, Cyp2a5 and, Hes1 shows their spatial expression patterns across all VNO cell types. Arrow highlights their specific expression in Sustentacular cells. **E)** Feature plot of immature neuronal marker Gap43 shows its lack of expression in the Sustentacular population (arrow mark). **F)** Feature plot of downstream Notch target Hey1 shows its expression in the Sustentacular population.

**Supplementary figure 4:**
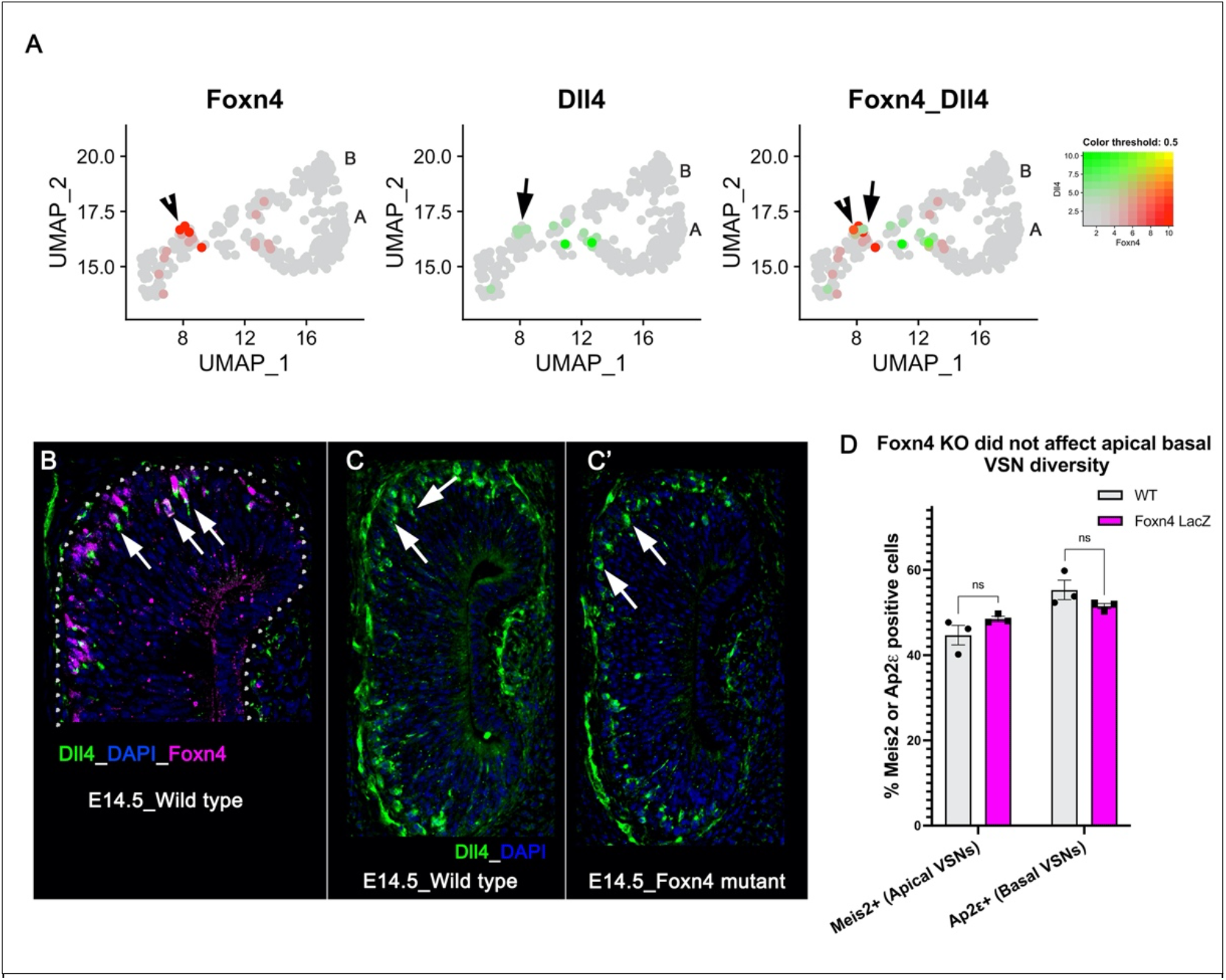
Foxn4 is redundant in inducing Dll4 expression. **A)** Blended feature plots of Foxn4 and Dll4 show that Foxn4 (arrowhead) is expressed at the dichotomy along with Dll4 (arrow). **B)** Double immunofluorescence of Dll4 and Foxn4 in wildtype VNO at embryonic day 14.5 (E14.5) stage shows their colocalization (arrow mark). **C, C’)** Immunofluorescence of Dll4 in wild type control and Foxn4 mutant VNOs show detectable levels of Dll4 expression (arrow marks). **D** Bar plot showing % Meis2 positive and Ap2ε positive cells in wild type and Foxn4 mutant VNOs at E14.5 stage

**Supplementary figure5:**
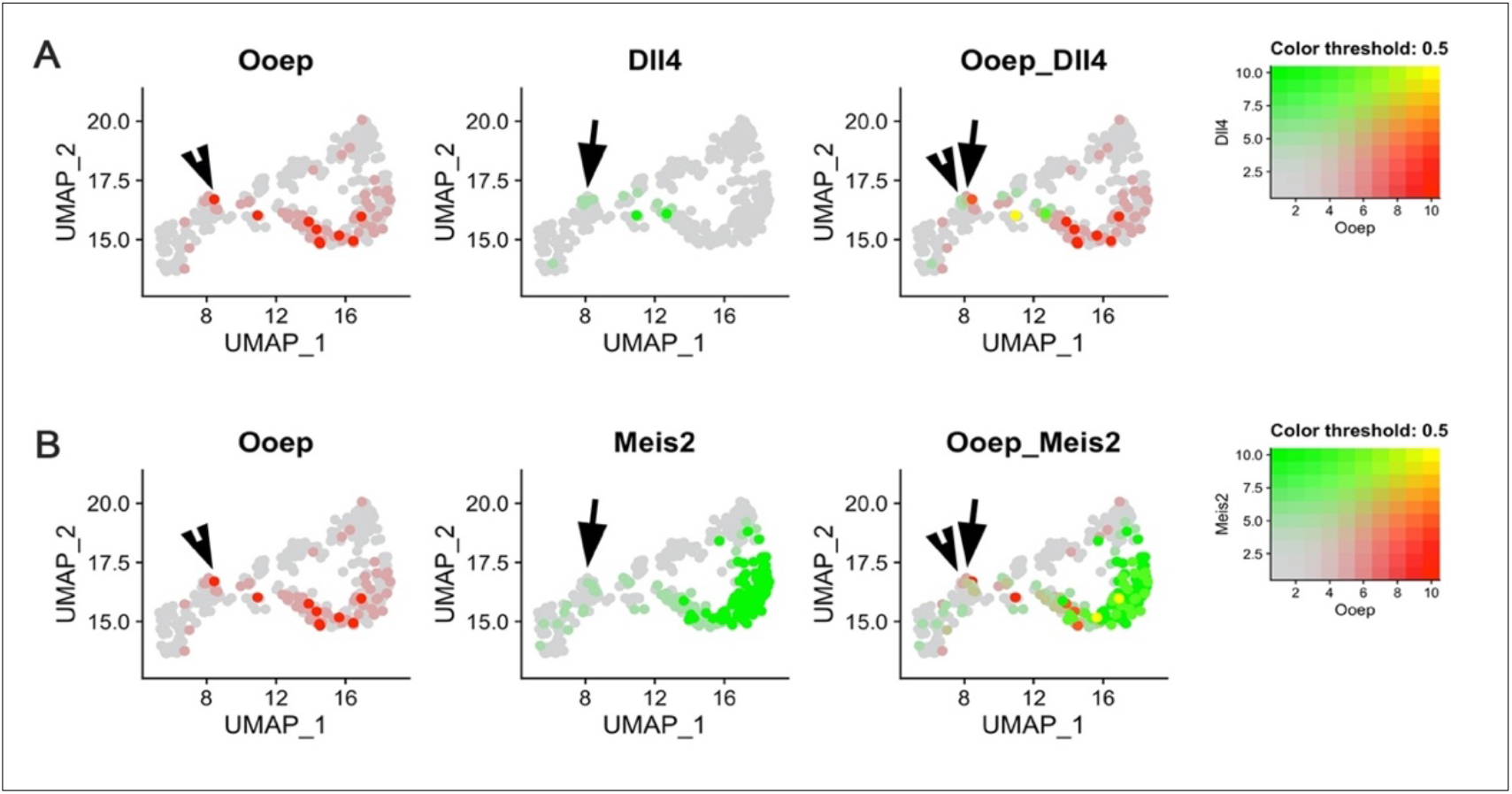
Oocyte expression protein (Ooep) is enriched in apical specific VSNs at the dichotomy. **A)** Blended feature plots of Ooep and Dll4 shows that Ooep (arrowhead) is initially expressed at the dichotomy, and it is enriched in Dll4 positive cells (arrow) **B)** Blended feature plots of Ooep and Meis2 shows that Ooep (arrowhead) is expressed along with apical VSN specific marker Meis2(arrow).

## Notes

### Competing Interest Statement

The authors have declared no competing interest.

